# Icosahedral viruses defined by their positively charged domains: a signature for viral identity and capsid assembly strategy

**DOI:** 10.1101/600981

**Authors:** Rodrigo D. Requião, Rodolfo L. Carneiro, Mariana Hoyer Moreira, Marcelo Ribeiro-Alves, Silvana Rossetto, Fernando L. Palhano, Tatiana Domitrovic

## Abstract

Capsid proteins often present a positively charged arginine-rich region at the N and/or C-termini that for some icosahedral viruses has a fundamental role in genome packaging and particle stability. These sequences show little to no conservation at the amino-acid level and are structurally dynamic so that they cannot be easily detected by common sequence or structure comparison. As a result, the occurrence and distribution of positively charged protein domain across the viral and the overall protein universe are unknown. We developed a methodology based on the net charge calculation of discrete segments of the protein sequence that allows us to identify proteins containing amino-acid stretches with an extremely high net charge. We observed that among all organisms, icosahedral viruses are especially enriched in extremely positively charged segments (Q ≥ +17), with a distinctive bias towards arginine instead of lysine. We used viral particle structural data to calculate the total electrostatic charge derived from the most positively charged protein segment of capsid proteins and correlated these values with genome charge arising from the phosphates of each nucleotide. We obtained a positive correlation (r = 0.91, p-value < 0001) for a group of 17 viral families, corresponding to 40% of all families with icosahedral structures described so far. These data indicated that unrelated viruses with diverse genome types adopt a common underlying mechanism for capsid assembly and genome stabilization based on R-arms. Outliers from a linear fit pointed to families with alternative strategies of capsid assembly and genome packaging.

**Significance Statement:** Viruses can be characterized by the existence of a capsid, an intricate proteinaceous container that encases the viral genome. Therefore, capsid assembly and function are essential to viral replication. Here we specify virus families with diverse capsid structure and sequence, for each capsid packing capacity depends on a distinctive structural feature: a highly positively charged segment of amino acids residues, preferentially made of arginine. We also show that proteins with the same characteristics are rarely found in cellular proteins. Therefore, we identified a conserved viral functional element that can be used to infer capsid assembly mechanisms and inspire the design of protein nanoparticles and broad-spectrum antiviral treatments.

## Introduction

The most common solution that viruses employ to protect their genomes is to assemble a spherical shell composed of multiple copies of only one or a few kinds of proteins. The capsid proteins (CP) interact with the genome and each other, usually following the principles of icosahedral symmetry, where the number of subunits forming the capsid is given by the triangulation number (T) × 60. The second architecture is a helical arrangement of proteins (nucleocapsid proteins, NCP) that interact with the genome (1, 2). The mechanisms involved in the assembly of the protein shell and condensation of viral capsids genome often find direct applications in the fields of drug development and nanotechnology.

Some icosahedral viruses have a high concentration of positively charged amino acid residues at the extremities of their CPs, known as arginine-rich motifs, poly-arginine or arginine-arms. These R-arms are directed towards the interior of the viral particle, where they make contact with the encapsulated nucleic acid (3). Studies with hepatitis B virus (4), circovirus (5), nodavirus (6), and other models (7, 8) have demonstrated that these positively charged domains are essential for interaction with the viral genome and for particle stability. Part of the functional explanation may rely on the counteraction of repulsive forces that results from the negatively charged nucleic acids condensed inside the capsid (9, 10). Different groups working with single-stranded positive sense (+) RNA viruses observed that the sum of net charges of all R-arm containing proteins in a virus capsid correlates with its genome packing capacity, e.g. (10–14). However, for some specific viruses, R-arms have also been implicated in the interaction with cellular membranes promoting particle penetration into the cell (15) or intracellular localization (16, 17). In these cases, R-arms can act as localization signals or cell penetrating peptides (18, 19), suggesting that these domains are multi-functional. Despite the notion that R-arms are present in different viruses and are critical components for viral replication and assembly, they have never been formally annotated as a protein domain by widely known resources and databases such as the Pfam protein family database (20) or InterPro (21). Consequently, there is no information on the distribution of R-arms across different organisms or viral families or their overall amino-acid composition. This broad view perspective is necessary to determine if R-arms can be considered a typical functional module of icosahedral viral capsids and if they can be used to infer capsid assembly mechanisms.

R-arms often present low sequence conservation and extensive variation in length, what hampers domain identification by profile Hidden Markov-Model (HMM) protein classification, the method employed by relevant databases such as Pfam (20). Moreover, R-arms often lie within an intrinsically disordered region, too dynamic or flexible to be resolved in viral capsid structural models generated by X-ray crystallography or cryo-electron microscopy. These attributes complicate the use of traditional approaches for the identification of R-arms in unrelated viruses; and sometimes, even within a viral family.

Here, to determine the occurrence of positively charged domains among protein from diverse organisms, including viruses, we analyzed the net charge distribution across the primary structure of proteins deposited in the reviewed Swiss-Prot database (559,052 proteins). Using a program that calculates the net charge in consecutive amino-acid stretches, we observed that icosahedral viruses are enriched with positively charged stretches when compared to *Homo sapiens* and other proteomes, especially at extreme charge values (≥ +17). The viral capsid segments also present at least 4 times more arginine than lysine, a feature that is not common in cellular proteins. We also made a focused effort to calculate the correlation between the total net-charge derived from the positively charged domain and the genome charge for a comprehensive group of viruses with different genome types. We propose that this analysis can be used to predict if the electrostatic interaction between the positively charged domain and the genome is a dominant driving force for capsid assembly and stability.

## Results

### The viral proteome contains more super-positive stretches than the cellular proteome

The first step to characterize the charge distribution along different protein sequences was to define the length of the search frame, that is, the number of residues that would be used for net-charge calculation in every consecutive stretch. Virus positively charged motifs can be in rigid patches on the inner capsid surface (e.g., bacteriophage MS2, *Leviviridae* family (22)) but usually they are within flexible arms in the N terminus. For this reason, a commonly used criterion for R-arm size determination is the length of the disordered region of the N-terminus as determined by x-ray crystal models (11) or secondary structure prediction software (23). We listed (+)RNA viruses that have been previously analyzed (10, 14, 23), and noticed that the average unstructured N-terminus is around 30 amino-acid residues (n= 14 families, SD ± 23.71). These observations indicated that this frame was a good starting point for our analysis.

In order to characterize the distribution of positively charged protein stretches in several organisms, we used a program that can screen a protein sequence and calculate the net charge every consecutive frame of 30-amino-acid residues (*Q*_30res_) (24). We analyzed total Swiss-Prot reviewed proteome (559,052 sequences) and compared with viral, *Drosophila melanogaster*, *Arabidopsis thaliana*, and *H. sapiens* data sets. In Fig. 1A, we show the frequency distribution of the stretches according to their *Q*_30res_. Different from insect, plant, and human, viruses had more positively charged than negatively charged segments. In Fig. 1B we show the log (base 10) net-charge frequency value for a selected group of proteins (e.g., Viral proteins) subtracted of the log expected frequency value calculated from the total proteome distribution (i.e., log10-fold-change). Viral proteins were the only class enriched in extremely high positively charged segments (charge ≥ + 17) when compared to the overall proteome (1B-inset with p-values). Positively charged protein stretches can be involved in diverse roles, such as membrane interaction, DNA or RNA binding, and cellular localization signaling (25). All these functions are important for virus replication and must contribute to the charge distribution profile of the viral protein dataset. In order to characterize the charge distribution according to protein function, we grouped the viral proteins following their functional annotation available in the Swiss-Prot database (Fig. 2). As expected, proteins classified in the DNA/RNA binding functional class (i.e., viral transcriptional factors, RNAi suppressors) had more positively charged segments than the Total Swiss-Prot proteome (Fig. 2A). However, the “Viral Particle” subset had higher frequencies of positively charged fragments (Fig. 2A). In Fig. 2B we dissected the viral particle components and observed that the class containing the highest frequencies and broadest distribution of positively charged segments was “Viral icosahedral capsid”. Even when compared to human DNA/RNA binding proteins, viral icosahedral capsid proteins concentrated more positively charged segments than any other analyzed classes (Fig. 2C).

**Figure 1.**
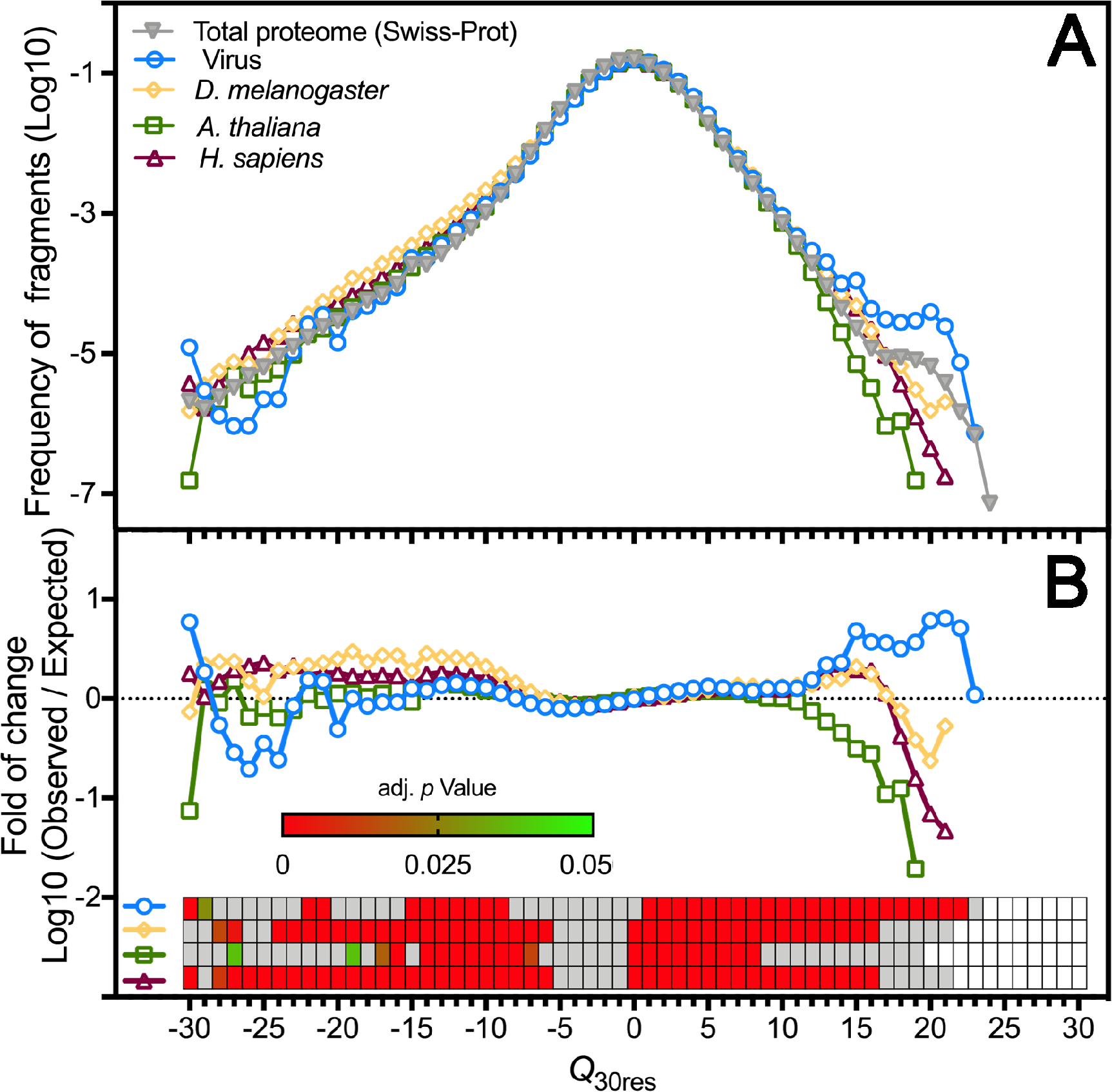
Viral proteins are enriched with positively charged stretches. Protein sequences derived from the reviewed Swiss-Prot data-bank were used as input for a program that calculates the net charge of every consecutive 30 amino-acid residues (*Q*_30res_). **A.** Normalized net-charge frequency distribution of protein segments from different organisms. **B.** Observed vs. expected net-charge frequency of each protein groups shown in A in relation to the Swiss-Prot proteome. The enrichment statistical analysis is shown in a heatmap (inset), where significant p-values are shown in shades of green to red. Grey slots represent p-values > 0.05.

**Figure 2.**
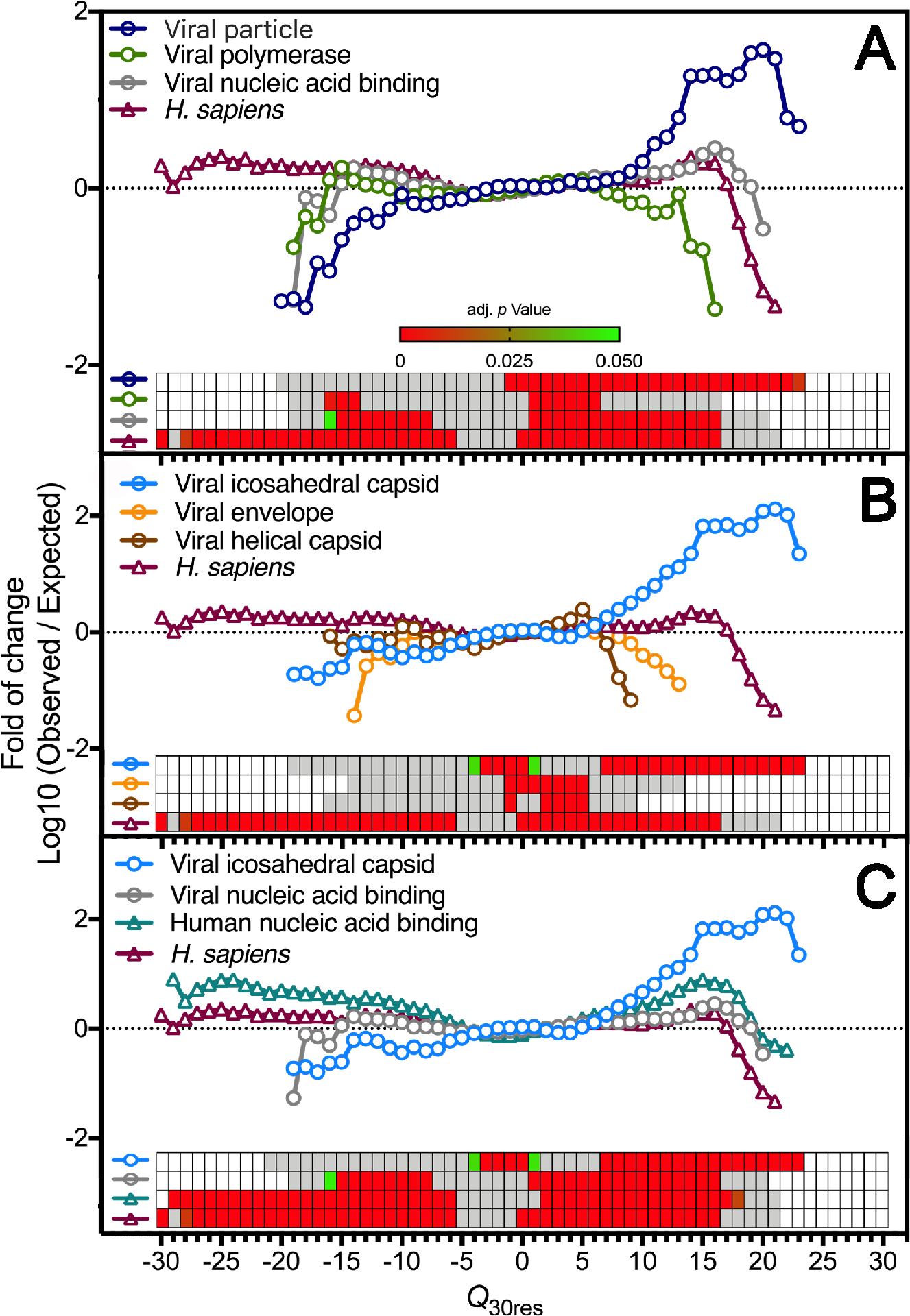
Capsid proteins from icosahedral viruses concentrate most of the positively charged protein segments of the viral proteome. Protein sequences derived from the reviewed Swiss-Prot data bank were used as input for a program that calculates the net charge of every consecutive 30 amino-acid residues (*Q*_30res_). The observed vs. expected frequency of fragments net charge from specific protein functional class, in relation to the Swiss-Prot proteome. **A.** The viral protein data set were divided into three different functional categories: Viral polymerase (containing all different kinds of viral polymerases); Nucleic acid binding (containing viral transcriptional/translational regulators, RNAi suppressors); and Viral particle (containing structural proteins present in viral particles). **B.** The Viral particle data-set was further divided into three different functional categories: Viral envelope (containing mainly glycoproteins); Viral helical capsid (containing mainly nucleocapsid proteins from helical viruses); and Viral icosahedral capsid (containing mainly capsid proteins from spherical viruses). **C.** The Viral icosahedral capsid data-set and the Viral nucleic acid binding dataset was compared to the Human nucleic acid binding data-set (containing RNA and DNA binding proteins with diverse functional roles). The statistical enrichment analysis is shown by a heatmap, where significant p-values are represented in shades of green to red. Grey slots represent p-values > 0.05.

### Positively charged domains of the icosahedral capsids are mainly involved in capsid assembly and stability

We hypothesized that by searching for the most positively charged segment in a capsid protein, we could efficiently identify the viral R-arm domains. Because the correlation between total R-arm charge and genome charge have already been demonstrated for a selected group of icosahedral RNA viruses (10, 14, 23), we decided to generalize this idea for all the icosahedral viruses in our dataset. This would not only validate our R-arm identification method for the previously analyzed (+)RNA viruses but would also reveal how the positively charged domain of viruses with different genome types relates to the capsid packaging capacity. While the theoretical determination of the genome charge (Q_*genome*_) is straightforward (each phosphodiester bond produces one negatively charged phosphate group), the calculation of R-arm total charge is more complicated. We carefully curated our protein dataset in order to select entries that corresponded to viruses with known capsid structure and complete genome sequence. The total R-arm net charge was calculated by multiplying the maximum net charge value in 30 amino-acid residues found in a protein capsid by the number of subunits forming the capsid (Total *Q*_max30res_). We accounted for deviations in icosahedral symmetry by using the actual subunit copy number (e.g., *Papillomaviridae*: pseudo T=7, with 72 pentamers of L1 and 72 copies of L2; *Geminiviridae*: formed by two fused T=1 capsids totalizing 110 subunits; *Picornaviridae*: pseudo T=3, with 60 copies of VP1, VP2, VP3, and VP4). We excluded viruses with complex multicomponent capsids; viruses with complex maturation pathways that involve scaffold proteins; and with uncertain protein copy number per capsid. With these criteria, we eliminated most bacteriophages (except *Leviviridae*) and complex icosahedral viruses, such as *Adenoviridae*, *Reoviridae*, *Herpesviridae,* etc. The final list (Support Table S1) contained 129 icosahedral viruses from 25 different families and all genome types, except for single-stranded negative sense (−)RNA (all helical viruses) and ssRNA-RT, comprising 57% of icosahedral virus families with known capsid structure (Viperdb). A linear fit allowing outliers identification indicated that 20 viruses, members of 8 virus families (marked in grey), deviated from the linear fit (Fig. 3A and Support Fig S2). Assuming these families as outliers, we analyzed the remaining 103 inliers from 17 families in a correlation analysis. We obtained a Pearson (r) of 0.91 and a p-value < 0.0001. In order to test the effect of different search frames, we repeated the analysis with segments of 10 and 60 amino-acid residues (Support Fig S2). The best linear fit was obtained with the 30 amino-acid residues frame (Total *Q*_max30res_ R^2^ = 0.74; Total *Q*_max10res_ R^2^ =0.42; Total *Q*_max60res_ R^2^ = 0.69). The fits from 30 and 60 aminoa-acid residues frames presented similar slopes (Total *Q*_max30res_ and Total *Q*_max60res_ slopes = 0.66 compared to *Q*_max10res_ slope = 0.42), and members from the same families were identified as outliers (S2). Therefore, we concluded that the 30-amino-acid residues frame was effectively reporting positively charged domains involved in genome stabilization, while the 10 amino-acid residue frame was too short to capture the entire positively charged domain. Figures 3B and C shows the *Q*_max30res_ and the *Q*_genome_ vs. T, respectively. Interestingly, T=1 ssDNA viruses carried segments with the highest net charge (Fig. 3B), probably to maximize packaging capacity of the smallest capsid of the viral world. Members from *Anelloviridae* and *Circoviridae* families have genome sizes equivalent to the geminated (T=1*) capsids of *Geminiviridae* (110 subunits), and even to some T=3 viruses, such as *Bromoviridae* (see Fig. 3A and 3B and supplemental table 1). We conclude that R-arms are a general strategy for capsid assembly and stabilization. Moreover, the identification of outliers indicates alternative functions for these domains and points to assembly strategies less dependent on electrostatic interactions between the genome and the capsid protein (see Discussion).

**Figure 3.**
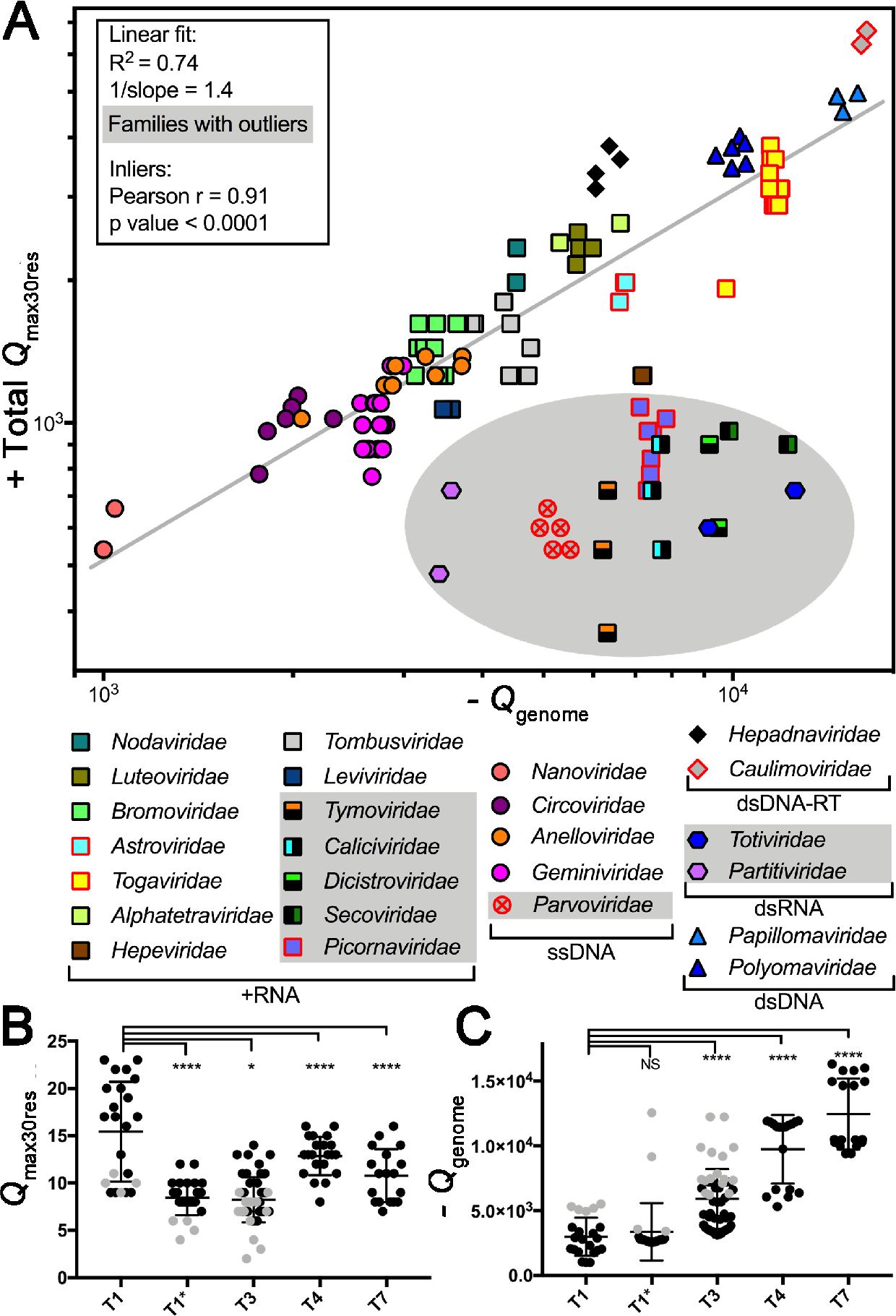
Total capsid internal net charge calculated from the most positively charged capsid protein segment correlates with genome packing capacity. **A.** The maximum net-charge value found in a 30 amino-acid residue stretch was multiplied by the number of subunits forming the capsid (Total *Q*_max30res_) of 129 viruses from 25 different families (See details in Support Table S1). The total nucleic-acid net charge was calculated from the number of nucleotides residues in the genome (Q_genome_). For multipartite viruses, the longest genome segment was considered for the plot. A straight line fit for the entire dataset was calculated, and a shaded area indicates the outliers (ROUT 2%). Pearson correlation results obtained from the inliers are shown in the inset. Data points contoured in red represent viruses that have more Lys than Arg in their positively charged segments (See also Fig 4A). **B** and **C** show the *Q*_max30res_ values per protein fragment and *Q*_genome_ values according to capsid T number, respectively. T1* corresponds to the T1 geminated capsids from *Geminiviridae* (110 subunits) and the dsRNA T1 capsids formed by dimeric subunits (120 subunits). Grey data points in B and C correspond to the outliers identified in panel A, as follows T1: ssDNA *Parvoviridae*; T1*: dsRNA *Totiviridae* and *Partitiviridae;* T3: all (+)RNA outliers *Caliciviridae*, *Dicistroviridae*, *Secoviridae, Picornaviridae* and *Tymoviridae*. Error bars indicate the mean and SD values. Tukey’s p-values **** < 0.0001, * 0.035.

Next, we examined the composition and location of these positively charged protein segments in viral capsid proteins (Fig 4). We complemented the protein dataset analyzed in Fig 3 (Fig 4, group 1 and 2) with icosahedral viruses with complex capsids (group 3) and helical viruses NCPs, totalizing 1,100 entries from 49 virus families. In Fig. 4A we show the ratio between arginine and lysine according to positive values of *Q*_30res_ found in CPs from different viral families. While around 60% of the viral proteins have more Arg than Lys residues in their most charged segment, the human proteome follows an opposite trend (see bottom Fig 4A). The bias towards arginine was stronger among the families that are included in the linear fit shown in Fig. 3A (see upper Fig 4A, group 1), but there were important exceptions. The inliers *Togaviridae* and *Caulimoviridae* have Lys-rich segments stabilizing their genomes. Among the helical viruses and the other icosahedral capsid proteins, we observed mixed patterns of R/K usage, but arginine is still preferred, especially in highly positively charged segments, such as the ones present in the histone-like proteins of adenoviruses (group 3). Another pattern that emerges from Fig 4A is that all viruses from group 1 have at least one segment with *Q*_30res_ ≥ +7. Even though this is not an exclusive feature of Group 1, we used this value as a threshold to map the location of positively charged segments in the primary structure of capsid proteins (Fig 4B). In order to allow a direct comparison, protein lengths were normalized and split into bins of 0.01; colors indicate the frequency of fragments with *Q*_30res_ ≥ +7. Helical viruses presented a more scattered and fragmented pattern of charge distribution than the inliers, which tend to have their positively charged segments concentrated in one or both extremities of the capsid protein, usually in the N-terminus. Finally, in Fig. 4C, we analyzed the amino-acid composition of the most charged segment of each virus from Group 1 *Q*_max30res_ and compared this with a dataset of *Q*_max30res_ of human nucleic-acid-binding proteins. Viruses had more arginine, proline, and tryptophan than the human dataset (Fig 4C). We looked for recurring patterns or known motifs in these sequences using MEME (data not shown) (26). The program retrieved expected motifs for the human data sets, such as RGG and RGR motifs for the human RNA-binding proteins (27); and zinc fingers and Homeobox motifs for the DNA-binding proteins (28). However, for the viral data set, no known nucleic-acid motifs were identified, and the few patterns retrieved by the program matched entries from the same family (not shown). This result confirms the unique structural makeup of viral capsid positively charged domains in relation to other DNA-and RNA-binding proteins.

**Figure 4.**
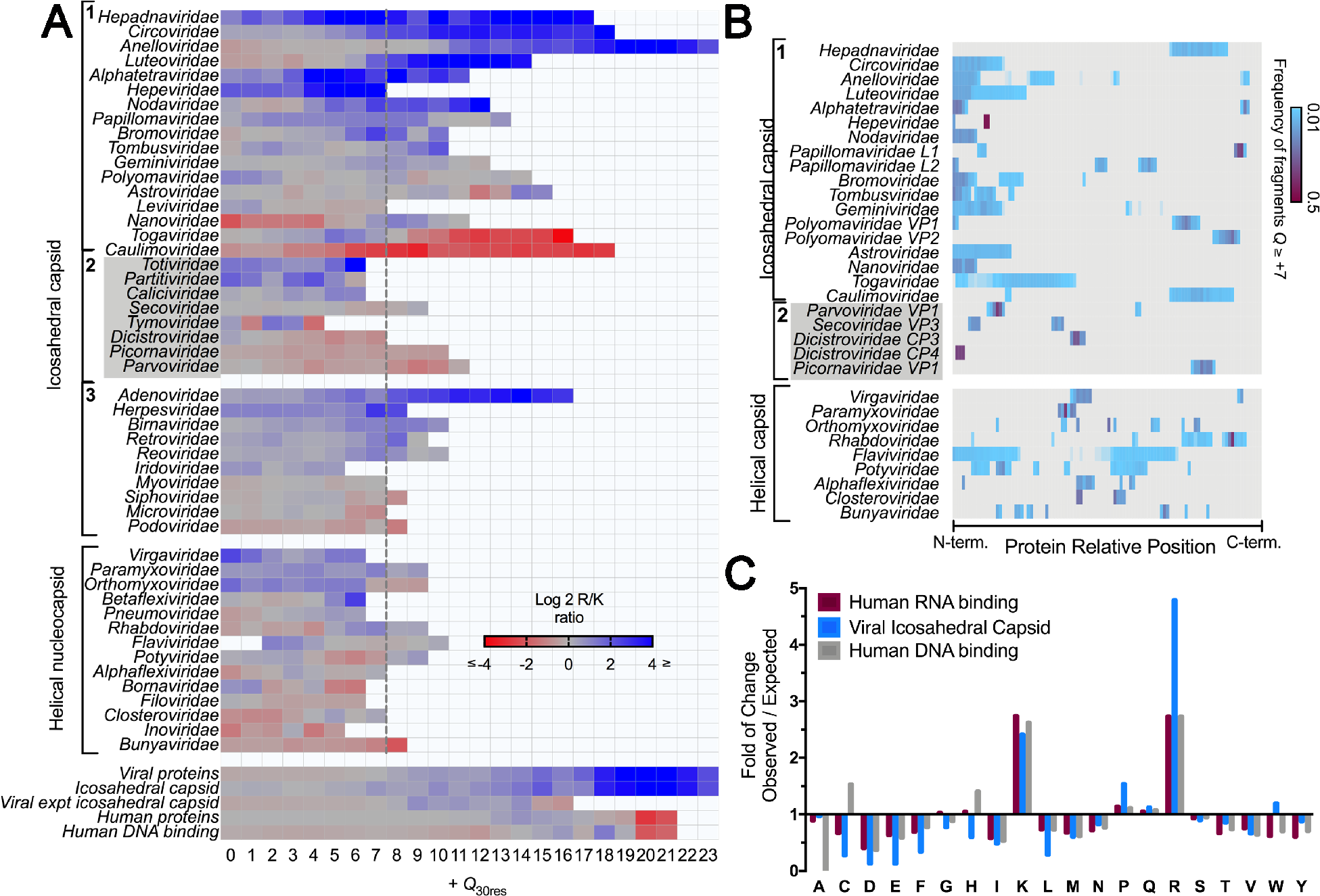
Composition and location of positively charged domains from viral capsid proteins: **A.** The arginine and lysine residues of fragments with *Q*_30res_ ≥ 0 of 1,100 capsid proteins from 49 virus families were calculated. The log (base = 2) R/K ratio per net charge values is shown as a heatmap, ranging from red (K-enriched) to blue (R-enriched). Groups 1 and 2 (grey box) contain the icosahedral viruses shown in Fig 3A; the latter corresponds to families identified as outliers. Group 3 contains complex multicomponent capsids that were not analyzed in Fig 3. From this plot, we see that all groups included in the linear fit of Figure 3A had at least one segment with Q_30res_ ≥ +7 (dashed line) **B.** A heatmap indicates the frequency values of fragments with Q_30res_ ≥ +7 according to their position in the primary structure. The proteins lengths were normalized and divided into bins of 0.01. **C.** The sequence of fragments with Q_max30res_ ≥ +7 from viral capsid proteins (group 1 panel A) and human nucleic acid binding proteins were used to determine the amino acid composition of positively charged segments. The panel shows the amino-acid enrichment in relation to the total Swiss-Prot proteome amino-acid composition.

## Discussion

We found that the high frequency of positively charged domains found in viruses (Fig 1) is due to the existence of icosahedral viral capsids (Fig 2), an extremely specialized quaternary arrangement of proteins and nucleic acids, which function and structure have no counterpart in cellular organisms. Only 0.1 % of all proteins of the Swiss-Prot database have at least one or more stretches with *Q*_30res_ ≥ +14 and R/K ≥ +4. About 25% of these are viral capsid proteins, a striking feature of viruses, considering that they represent only 3% of the Swiss-Prot proteome (Support Fig S3). Vertebrates are the second group having a considerable number of proteins with a similar constitution. Nevertheless, these proteins represent a tiny fraction of the individual organisms proteome (e.g., just 55 proteins with Q_30res_ ≥ +14 and R/K ≥ 4 in 20,415 human proteins). Among them, nucleic acid binding proteins, and more notably, protamines, which are small proteins expressed exclusively during spermatogenesis and involved in DNA hyper-condensation (29) (Support Fig S3). The arginine side chain possesses a guanidinium group, able to form bidentate bonds that are advantageous to maximize nucleic acid folding and packing when compared to Lys (30, 31). Moreover, arginine-rich cell-penetrating peptides are more efficient than the lysine-rich peptides, probably because of the bidentate interaction forces membrane curvature and destabilization (32). Hence, arginine seems to be the optimal amino acid to condense and stabilize the viral genome and to facilitate membrane interaction. Nevertheless, different from the negatively charged amino acids, the concentration of R and K in a short protein segment is limited (Fig. 1). The adverse effect of exceptionally positively charged protein segments on ribosomal synthesis efficiency may be among the selective pressures acting against repetitions of R or K in all organisms (24). Additionally, the size and composition of viral positively charged domains might be controlled by other factors. Viral nucleic-acid structural features that are rare in host cells usually serve as molecular targets for innate immune response (33), and is possible that R-rich domains function as a viral protein specific pattern.

The calculation of capsid internal net charge shown in Figure 3 follows the most straightforward methodology published so far (11, 14, 23) since the only criterion for R-arm identification is the assumption that it is the most positively charged segment of the capsid protein. Even so, we observed a positive correlation between total *Q*_max30res_ and *Q*_genome_ for all the (+)RNA viruses for each the involvement of positively charged domains and capsid assembly was experimentally demonstrated (e.g., *Geminiviridae*, *Bromoviridae*, *Togaviridae*). The 1/slope value that gives the capsid/genome charge ratio was 1.4, which is generally in line with previous data indicating that (+)RNA viral capsids are overcharged, meaning that the *Q*_genome_ is not completely neutralized by protein derived positive charges (3, 9, 10). Despite the simplicity of the calculation, we reproduced the general findings obtained with (+)RNA viruses (3) (23) and showed that the striking correlation between genome charge and positively charged protein domains could actually be extended to other groups of viruses with different genome types. While our analysis implies a general role for positively charged domains in capsid assembly and genome interaction for all the families included in the linear fit, is important to note that the assembly pathways, and the functional details can be quite diversified. Some viral capsids rely more heavily on CP-CP interactions for assembly, as suggested by the formation of empty capsids in the absence of positively charged domains (e.g., *Hepadnaviridae* (4)), while others are entirely dependent on R-arms to form the capsid (e.g., *Nodaviridae* (16) and *Alphatetraviridae* (John E. Johnson, personal communication). *Polyomaviridae* and *Papillomaviridae* are known to pack their genome with histones, suggesting that the R-arms are not sufficient to stabilize or condense the stiffer dsDNA (34). The 8 outliers viruses families in the linear fit represent alternative assembly strategies with little contribution of electrostatic interactions between the capsid and the genome. In all cases, the genome charge exceeded the expected internal capsid charge (Fig. 3A). These groups contained the two dsRNA virus families included in the plot, as well as 6 ssRNA and one ssDNA virus family.

The dsRNA families *Totiviridae* and *Partitiviridae* share similar simple capsid architecture, with 60 CP dimers forming a T1 capsid. All dsRNA viruses, including the more complex reoviruses and birnaviruses, replicate their genome and transcribe their mRNA inside an assembled capsid that also encloses the RNA-dependent RNA polymerase. More than transporting the genome, these particles are part of the viral factory, preventing the detection of viral dsRNA species by cellular proteins (35). Because these capsids must sustain variable levels of RNA content during viral replication, it is reasonable that these families diverged from the group belonging to the linear fit. Among the ssRNA outliers, we found *Caliciviridae*; the 3 families of picornavirales present in the dataset (*Dicistroviridae*, *Secoviridae*, and *Picornaviridae*); and *Tymoviridae*. A recent sequence-similarity network analysis of single Jelly-Roll capsid proteins from RNA viruses revealed two large clusters, one containing most of the ssRNA viruses present in our data-set and another formed by picornavirales and *Caliciviridae (36)*. Even though the capsid architecture is not the same, both groups pack VPg, a small protein bound to the genome 5’-end (37). Picornaviruses form pseudo-T = 3 viruses containing 4 different proteins. Segments with charge > +7 were found in few entries and were restricted to one or two CPs. The primary role of these domains is unknown, but they may participate in membrane interaction, as already demonstrated for dicistroviruses CP4 (38). Most viruses from *Caliciviridae* assemble their capsid with one type of CP arranged in 90 dimers in a T=3 lattice. No segments with Q ≥ +7 were found in *Caliciviridae* CPs. Our data reinforce the structural similarities between these two groups and suggest a common, yet unknown mechanism for genome stabilization and assembly. The *Tymoviridae* capsid proteins are also devoided of segments with Q ≥ +7. An X-ray structural model of DYMV include densities corresponding to ordered RNA inside the capsid, but no positively charged residues are present in the interaction interface (39). The *Parvoviridae* were the only T = 1 ssDNA viruses identified as an outlier family. These viruses enclose the largest genomes among the ssDNA viruses (~ 5 kb) but have charge values similar to the tiny *Nanoviridae* (~ 1 kb). Parvoviruses present 3 variations of the cap gene product, all having an overlapping amino-acid sequence with similar C-termini. The most charged segment is a short Lys-enriched region unique to VP1. Because this CP variant is the least abundant, our charge calculation is probably overestimated. The capsid is mainly formed by VP2 proteins that have a very conserved ssDNA binding pocket (40). The binding site shows an ordered loop of 9 nucleotides that coordinates two Mg^2+^. This stable and structured contact between the genome and the protein shell may represent an alternative strategy to the long super-charged R-arms that are observed in circovirus and anellovirus (5, 40). ssDNA viruses are known to be a polyphyletic diversified group (41). The finding that *Parvoviridae* has genome stabilization strategy that differs from other small ssDNA viruses only contributes to the hypothesis of independent events of capsid acquisition in this group.

In summary, positively charged domains that are implicated in viral capsid stabilization have general features such as, possessing a high positive charge in a 30 amino-acid residue stretch (*Q*_30res_ ≥ +7); being enriched in arginine over lysine; and being located at the C- or N-terminus of the capsid protein. However, these characteristics are neither essential nor exclusive to genome stabilization function, which complicates a sequence only approach to R-arms identification. On the other hand, when associated with virus genome charge and capsid structure, positively charged domains can suggest the general basis of capsid assembly and genome packaging mechanisms.

## Methodology

### Data sources

Protein database Swiss-Prot at Uniprot.org was used as our source of primary protein sequences. Protein function, taxonomic and structural information were retrieved from Uniprot.org, Viralzone, and Viperdb. Genome sizes for all viruses were obtained at the National Center for Biotechnology Information (NCBI) database. Reference sequences were used when available.

### Net charge calculation, R/K ratios determination, amino-acid composition, and statistics

We developed a program that screens the primary sequence of a given protein and calculates the net charge in consecutive frames of a predetermined number of amino acids (10, 30 or 60 were used). For the net charge determination, K and R were considered +1; D and E were considered −1; every other residue was considered 0. The N and C termini charges were disregarded. In a previous publication, we have shown that these simplified parameters are equivalent to a calculation using partial charges of individual amino acids at pH 7.4, according to their p*K*a values and Henderson-Hasselbach equation (24). After net charge calculation in 30 amino-acid residue stretches, another program was used to determine the arginine (R) to lysine (K) ratio in a group of amino-acid stretches using the equation available in the support information S4. The general amino-acid composition of 30 residues stretches with *Q* ≥ +7 was generated by MEME (http://meme-suite.org) (26). Statistical analyses and graphical displays were made either with R version 3.5.2 or Graph-pad Prism 7.0 software. The details of *in silico* implementation and statistics are described in Support Information and the code software’s were posted at https://github.com (see also Support Information S4).

## Supporting information

Support Information

## References

1. Caspar DL & Klug A (1962) Physical principles in the construction of regular viruses. Cold Spring Harbor symposia on quantitative biology 27:1–24.

2. Johnson JE & Speir JA (1997) Quasi-equivalent viruses: a paradigm for protein assemblies. Journal of molecular biology 269(5):665–675.

3. Perlmutter JD & Hagan MF (2015) Mechanisms of virus assembly. Annual review of physical chemistry 66:217–239.

4. Newman M, Chua PK, Tang FM, Su PY, & Shih C (2009) Testing an electrostatic interaction hypothesis of hepatitis B virus capsid stability by using an in vitro capsid disassembly/reassembly system. Journal of virology 83(20):10616–10626.

5. Khayat R, et al. (2011) The 2.3-angstrom structure of porcine circovirus 2. Journal of virology 85(15):7856–7862.

6. Schneemann A & Marshall D (1998) Specific encapsidation of nodavirus RNAs is mediated through the C terminus of capsid precursor protein alpha. Journal of virology 72(11):8738–8746.

7. Garmann RF, Comas-Garcia M, Gopal A, Knobler CM, & Gelbart WM (2014) The assembly pathway of an icosahedral single-stranded RNA virus depends on the strength of inter-subunit attractions. Journal of molecular biology 426(5):1050–1060.

8. Rayaprolu V, et al. (2017) Length of encapsidated cargo impacts stability and structure of in vitro assembled alphavirus core-like particles. J Phys Condens Matter 29(48):484003.

9. Garmann RF, Comas-Garcia M, Knobler CM, & Gelbart WM (2016) Physical Principles in the Self-Assembly of a Simple Spherical Virus. Acc Chem Res 49(1):48–55.

10. Perlmutter JD, Qiao C, & Hagan MF (2013) Viral genome structures are optimal for capsid assembly. Elife 2:e00632.

11. Belyi VA & Muthukumar M (2006) Electrostatic origin of the genome packing in viruses. Proceedings of the National Academy of Sciences of the United States of America 103(46):17174–17178.

12. Siber A & Podgornik R (2008) Nonspecific interactions in spontaneous assembly of empty versus functional single-stranded RNA viruses. Physical review. E, Statistical, nonlinear, and soft matter physics 78(5 Pt 1):051915.

13. Ting CL, Wu J, & Wang ZG (2011) Thermodynamic basis for the genome to capsid charge relationship in viral encapsidation. Proceedings of the National Academy of Sciences of the United States of America 108(41):16986–16991.

14. Hu TZ R.; Shklovskii, B. I. (2008) Electrostatic theory viral self-assembly. Physica A 387(12):3059–3064.

15. Domitrovic T, Matsui T, & Johnson JE (2012) Dissecting Quasi-Equivalence in Nonenveloped Viruses: Membrane Disruption Is Promoted by Lytic Peptides Released from Subunit Pentamers, Not Hexamers. Journal of virology 86(18):9976–9982.

16. Venter PA, Marshall D, & Schneemann A (2009) Dual roles for an arginine-rich motif in specific genome recognition and localization of viral coat protein to RNA replication sites in flock house virus-infected cells. Journal of virology 83(7):2872–2882.

17. Sarker S, et al. (2016) Structural insights into the assembly and regulation of distinct viral capsid complexes. Nat Commun 7:13014.

18. Freire JM, et al. (2013) Intracellular nucleic acid delivery by the supercharged dengue virus capsid protein. PloS one 8(12):e81450.

19. Zhang P, Monteiro da Silva G, Deatherage C, Burd C, & DiMaio D (2018) Cell-Penetrating Peptide Mediates Intracellular Membrane Passage of Human Papillomavirus L2 Protein to Trigger Retrograde Trafficking. Cell 174(6):1465–1476 e1413.

20. Finn RD, et al. (2016) The Pfam protein families database: towards a more sustainable future. Nucleic acids research 44(D1):D279–285.

21. Richardson LJ, et al. (2018) Genome properties in 2019: a new companion database to InterPro for the inference of complete functional attributes. Nucleic acids research.

22. Valegard K, et al. (1997) The three-dimensional structures of two complexes between recombinant MS2 capsids and RNA operator fragments reveal sequence-specific protein-RNA interactions. Journal of molecular biology 270(5):724–738.

23. Losdorfer Bozic A & Podgornik R (2018) Varieties of charge distributions in coat proteins of ssRNA+ viruses. J Phys Condens Matter 30(2):024001.

24. Requiao RD, et al. (2017) Protein charge distribution in proteomes and its impact on translation. PLoS Comput Biol 13(5):e1005549.

25. Garcia-Moreno M, Jarvelin AI, & Castello A (2018) Unconventional RNA-binding proteins step into the virus-host battlefront. Wiley Interdiscip Rev RNA 9(6):e1498.

26. Bailey TL, Johnson J, Grant CE, & Noble WS (2015) The MEME Suite. Nucleic acids research 43(W1):W39–49.

27. Hentze MW, Castello A, Schwarzl T, & Preiss T (2018) A brave new world of RNA-binding proteins. Nat Rev Mol Cell Biol 19(5):327–341.

28. Johnson PF & McKnight SL (1989) Eukaryotic transcriptional regulatory proteins. Annual review of biochemistry 58:799–839.

29. Braun RE (2001) Packaging paternal chromosomes with protamine. Nature genetics 28(1):10–12.

30. DeRouchey J, Hoover B, & Rau DC (2013) A comparison of DNA compaction by arginine and lysine peptides: a physical basis for arginine rich protamines. Biochemistry 52(17):3000–3009.

31. Ni P, et al. (2012) An examination of the electrostatic interactions between the N-terminal tail of the Brome Mosaic Virus coat protein and encapsidated RNAs. Journal of molecular biology 419(5):284–300.

32. Kalafatovic D & Giralt E (2017) Cell-Penetrating Peptides: Design Strategies beyond Primary Structure and Amphipathicity. Molecules (Basel, Switzerland) 22(11).

33. Brennan K & Bowie AG (2010) Activation of host pattern recognition receptors by viruses. Current opinion in microbiology 13(4):503–507.

34. Hurdiss DL, et al. (2016) New Structural Insights into the Genome and Minor Capsid Proteins of BK Polyomavirus using Cryo-Electron Microscopy. Structure 24(4):528–536.

35. Luque D, Mata CP, Suzuki N, Ghabrial SA, & Caston JR (2018) Capsid Structure of dsRNA Fungal Viruses. Viruses 10(9).

36. Wolf YI, et al. (2018) Origins and Evolution of the Global RNA Virome. MBio 9(6).

37. Goodfellow I (2011) The genome-linked protein VPg of vertebrate viruses - a multifaceted protein. Current opinion in virology 1(5):355–362.

38. Sanchez-Eugenia R, Goikolea J, Gil-Carton D, Sanchez-Magraner L, & Guerin DM (2015) Triatoma virus recombinant VP4 protein induces membrane permeability through dynamic pores. Journal of virology 89(8):4645–4654.

39. Larson SB, Day J, Canady MA, Greenwood A, & McPherson A (2000) Refined structure of desmodium yellow mottle tymovirus at 2.7 A resolution. Journal of molecular biology 301(3):625–642.

40. Halder S, et al. (2013) Structural characterization of H-1 parvovirus: comparison of infectious virions to empty capsids. Journal of virology 87(9):5128–5140.

41. Krupovic M (2013) Networks of evolutionary interactions underlying the polyphyletic origin of ssDNA viruses. Current opinion in virology 3(5):578–586.

