## Supplementary material for "Icosahedral viruses defined by their positively charged domains: a signature for viral identity and capsid assembly strategy": Support Information

**Support Method Information for:**

#### MATERIALS AND METHODS

| Software and Algorithms | Source | Identifier |
|---|---|---|
| Calculation of net charge | (1) | |
| Calculation of R/K ratio | This paper | https://github.com/mhoyerm/Total_ratio |
| Identify proteins of determined net charge and R/K ratio | This paper | https://github.com/mhoyerm/Modulate_RK |
| Identify proteins of determined net charge and K/R ratio | This paper | https://github.com/mhoyerm/Modulate_KR |

Data sources

For all viral proteins, we used UniRef with the advanced search options (uniprot:(proteome:(taxonomy:"Viruses [10239]") reviewed:yes) AND identity:1.0). For viral capsid proteins, we used the advanced search options (proteome:(taxonomy:"Viruses [10239]") goa:("viral capsid [19028]") AND reviewed:yes) followed by a manual selection of major capsid proteins. Advanced search options for H. sapiens (reviewed:yes AND organism:"Homo sapiens (Human) [9606]"), D. melanogaster (reviewed:yes AND organism:"Drosophila melanogaster (Fruit fly) [7227]") and A. thaliana (reviewed:yes AND organism:"Arabidopsis thaliana (Mouse-ear cress) [3702]") were used as a source for the proteomes. For protein classes analyses, proteins were further separated by their Gene Ontology ID (GO:3677 DNA binding; GO:3723 RNA binding; GO:34061 DNA polymerase activity; GO:34062 5'-3' RNA polymerase activity; GO:19031 viral envelope). Viral capsid proteins were separated in helical or non-helical by helical viral capsid Gene Ontology ID (GO:19029).

Net charge calculation.

To calculate the net charge in a given sequence, a program that can screen the primary sequence of a given protein and attribute different values to different amino acids was developed. To determine the charge values for the amino-acid residues, we considered the contribution of individual pKa values to the Henderson-Hasselbach equation at pH 7.4 (1). The final peptide net-charge was determined by summing each amino-acid charge. The following charge values were obtained: lysine = +0.999 (rounded value = + 1); arginine = +1.000; histidine = +0.048 (rounded value = 0); glutamic acid = -0.999 (rounded value = -1); aspartic acid = -1.000; cysteine = -0.085 (rounded value = 0) and all other amino acids = 0.000. The N- and C-termini were attributed to the charges +0.996 and -1.000, respectively. After the charge attribution, the program calculated the net-charge of every 30-amino-acid frame in the primary sequence. Because the results obtained by the program that used non-rounded values were very similar to the rounded values (data not shown), in all calculations used herein, the rounded values were used.

Overrepresentation of net charge values varying from -30 to +30 calculated in 30-width amino-acid stretches were performed by right-sided P[X ≥x] hypergeometric tests. For the enrichment analysis of any specific net charge value, hypergeometric distributions were used for sampling without replacement. The density of these distributions with parameters *m*, *n* and *k* were given by Equation 1

$$p(x) = \frac{m}{n} \times \frac{n}{k-x} / \frac{m+n}{k}, \forall x = 0, \ldots, k, \qquad \text{Eq.1}$$

where, *x* was the number (-1) of amino-acid stretches in proteins of any specific species (e.g. *D. melanogaster*, *A. thaliana*, *H. sapiens*) or categories (e.g. Viral particle, polymerase, nucleic acid binding) with any specific net charge; *m* were the number of amino-acid stretches in all Swiss-prot database proteins with any specific net-charge; *n* was the number of all Swiss-prot database of any net-charge but the one being analyzed; and, *k* was the number of amino-acid stretches of any charge in proteins of any species or categories being analyzed. Accordingly, the expected number of amino-acid stretches in proteins of any specific species or categories with any net charge value were calculated by *m*×(*k* / (*m*+*n*).

Identification of proteins with positive net charge and biased towards lysines or arginines.

To study the identity of proteins with positive net charge stretches enriched on arginine's (R) or lysine's (K), we designed and implemented another algorithm that calculated the net charge of amino acids at a determined ratio of R/K or K/R. The calculations of net charge, total arginine, total lysine and the ratio among R/K or K/R were calculated by Equation 2.

$$J_j = \frac{\sum_{i=0}^{n} R_{ij}}{\sum_{i=0}^{n} K_{ij}}, \qquad \text{Eq.2}$$

where, *Jj* is ratio R/K for the respective *j* net charge, $R_{ij}$ is the number of arginines found in stretch *i*, $K_{ij}$ is the number of lysines found in stretch *i*, and n is the total number of stretches found.

S1: Supplemental
[illegible]

| | | | | | | Max Net charge | | | | Total net charge | | |
|---|---|---|---|---|---|---|---|---|---|---|---|---|
| Protein UniprotKB ID | Genome reference | Genome size | Family | Genome type | T | 10 aa | 30 aa | 60 aa | Genome charge | 10aa | 30aa | 60aa |
| P03089 | NC_001699.1 | 5130 | Polyomaviridae | dsDNA | 7 | 6 | 9 | 9 | 10260 | 2592 | 4032 | 3960 |
| P03095 | NC_001699.1 | 5130 | Polyomaviridae | dsDNA | 7 | 6 | 11 | 10 | | | | |
| A5HBD5 | NC_009539.1 | 5229 | Polyomaviridae | dsDNA | 7 | 5 | 8 | 10 | 10458 | 2304 | 3888 | 5184 |
| A5HBE1 | NC_009539.1 | 5229 | Polyomaviridae | dsDNA | 7 | 7 | 14 | 22 | | | | |
| P24848 | NC_001442.1 | 4697 | Polyomaviridae | dsDNA | 7 | 4 | 8 | 6 | 9394 | 1800 | 3672 | 2448 |
| P24849 | NC_001442.1 | 4697 | Polyomaviridae | dsDNA | 7 | 5 | 11 | 4 | | | | |
| P03087 | NC_001669.1 | 5243 | Polyomaviridae | dsDNA | 7 | 5 | 7 | 8 | 10486 | 2232 | 3528 | 3888 |
| P03093 | NC_001669.1 | 5243 | Polyomaviridae | dsDNA | 7 | 6 | 14 | 14 | | | | |
| P13891 | NC_004764.2 | 4981 | Polyomaviridae | dsDNA | 7 | 4 | 8 | 6 | 9962 | 1944 | 3816 | 3096 |
| P13892 | NC_004764.2 | 4981 | Polyomaviridae | dsDNA | 7 | 7 | 13 | 13 | | | | |
| P22163 | KT944080.1 | 7898 | Papillomaviridae | dsDNA | 7 | 7 | 12 | 11 | 15796 | 2952 | 4968 | 4608 |
| P22165 | KT944080.1 | 7898 | Papillomaviridae | dsDNA | 7 | 6 | 9 | 9 | | | | |
| P04012 | NC_027779.1 | 7320 | Papillomaviridae | dsDNA | 7 | 7 | 12 | 13 | 14640 | 3024 | 4896 | 5184 |
| P04013 | NC_027779.1 | 7320 | Papillomaviridae | dsDNA | 7 | 7 | 8 | 7 | | | | |
| P03103 | NC_035208.1 | 7480 | Papillomaviridae | dsDNA | 7 | 6 | 11 | 11 | 14960 | 2520 | 4536 | 4392 |
| Q705F9 | NC_035208.1 | 7480 | Papillomaviridae | dsDNA | 7 | 5 | 8 | 6 | | | | |
| Q66283 | NC_001648.1 | 8159 | Caulimoviridae | dsDNA-RT | 7 | 6 | 16 | 24 | 16318 | 2520 | 6720 | 10080 |
| P27502 | NC_001914.1 | 8002 | Caulimoviridae | dsDNA-RT | 7 | 6 | 15 | 19 | 16004 | 2520 | 6300 | 7980 |
| P0C677 | NC_003977.2 | 3182 | Hepadnaviridae | dsDNA-RT | 4 | 7 | 16 | 17 | 6364 | 1680 | 3840 | 4080 |
| P0C6J1 | NC_001484.1 | 3311 | Hepadnaviridae | dsDNA-RT | 4 | 6 | 15 | 17 | 6622 | 1440 | 3600 | 4080 |
| P0C6K0 | NC_001486.1 | 3027 | Hepadnaviridae | dsDNA-RT | 4 | 7 | 14 | 20 | 6054 | 1680 | 3360 | 4800 |
| Q66406 | NC_001344.1 | 3027 | Hepadnaviridae | dsDNA-RT | 4 | 6 | 13 | 18 | 6054 | 1440 | 3120 | 4320 |
| O15926 | U95995.1 | 1786 | Partitiviridae | dsRNA | 2 | 6 | 6 | 8 | 3572 | 720 | 720 | 960 |
| Q64FN9 | NC_006276.1 | 1708 | Partitiviridae | dsRNA | 2 | 4 | 4 | 5 | 3416 | 480 | 480 | 600 |
| Q67652 | NC_003555.1 | 6277 | Totiviridae | dsRNA | 2 | 4 | 6 | 8 | 12554 | 480 | 840 | 960 |
| P32503 | NC_003745.1 | 4579 | Totiviridae | dsRNA | 2 | 4 | 5 | 5 | 9158 | 480 | 600 | 600 |
| Q9DUB7 | NC_014087.1 | 3718 | Anelloviridae | ssDNA | 1 | 9 | 23 | 41 | 3718 | 540 | 1380 | 2460 |
| A4GZ97 | AB290918 | 3249 | Anelloviridae | ssDNA | 1 | 9 | 23 | 36 | 3249 | 540 | 1380 | 2160 |
| A7XCE4 | NC_015783.1 | 3709 | Anelloviridae | ssDNA | 1 | 8 | 22 | 41 | 3709 | 480 | 1320 | 2460 |
| Q9DUC1 | NC_014085.1 | 3371 | Anelloviridae | ssDNA | 1 | 9 | 21 | 38 | 3371 | 540 | 1260 | 2280 |
| Q8QVL6 | NC_014071.1 | 2797 | Anelloviridae | ssDNA | 1 | 8 | 20 | 31 | 2797 | 480 | 1200 | 1860 |
| Q9QU30 | NC_020498.1 | 2912 | Anelloviridae | ssDNA | 1 | 8 | 22 | 31 | 2912 | 480 | 1320 | 1860 |
| Q8QVL9 | AB076001 | 2878 | Anelloviridae | ssDNA | 1 | 8 | 20 | 35 | 2878 | 480 | 1200 | 2100 |
| Q8QVL3 | NC_014072.1 | 2064 | Anelloviridae | ssDNA | 1 | 8 | 17 | 28 | 2064 | 480 | 1020 | 1680 |
| Q9IG42 | NC_002361.1 | 2037 | Circoviridae | ssDNA | 1 | 8 | 19 | 26 | 2037 | 480 | 1140 | 1560 |
| Q9YUC8 | NC_001944.1 | 1993 | Circoviridae | ssDNA | 1 | 8 | 18 | 24 | 1993 | 480 | 1080 | 1440 |
| O56129 | NC_005148.1 | 1768 | Circoviridae | ssDNA | 1 | 8 | 13 | 22 | 1768 | 480 | 780 | 1320 |
| Q8QME6 | NC_003410.1 | 1949 | Circoviridae | ssDNA | 1 | 7 | 17 | 22 | 1949 | 420 | 1020 | 1320 |
| Q99153 | NC_001427.1 | 2319 | Circoviridae | ssDNA | 1 | 7 | 17 | 27 | 2319 | 420 | 1020 | 1620 |
| Q91EK1 | NC_003054.1 | 1821 | Circoviridae | ssDNA | 1 | 8 | 16 | 20 | 1821 | 480 | 960 | 1200 |
| P14984 | NC_001412.1 | 2993 | Geminiviridae | ssDNA | 1 | 5 | 12 | 16 | 2993 | 550 | 1320 | 1760 |
| Q88886 | NC_003825.1 | 2861 | Geminiviridae | ssDNA | 1 | 7 | 12 | 16 | 2861 | 770 | 1320 | 1760 |
| O39520 | NC_003493.2 | 2561 | Geminiviridae | ssDNA | 1 | 5 | 10 | 10 | 2561 | 550 | 1100 | 1100 |
| P03569 | NC_001346.2 | 2689 | Geminiviridae | ssDNA | 1 | 5 | 10 | 10 | 2689 | 550 | 1100 | 1100 |
| Q9YPS5 | NC_001983.1 (segment a) | 2723 | Geminiviridae | ssDNA | 1 | 5 | 10 | 13 | 2723 | 550 | 1100 | 1430 |
| Q00323 | NC_001647.1 | 2705 | Geminiviridae | ssDNA | 1 | 6 | 10 | 11 | 2705 | 660 | 1100 | 1210 |
| Q89551 | NC_003744.1 | 2758 | Geminiviridae | ssDNA | 1 | 5 | 10 | 11 | 2758 | 550 | 1100 | 1210 |
| P31616 | NC_003822.1 | 2580 | Geminiviridae | ssDNA | 1 | 5 | 9 | 10 | 2580 | 550 | 990 | 1100 |
| Q08583 | NC_001932.1 (segment 1) | 2815 | Geminiviridae | ssDNA | 1 | 5 | 9 | 14 | 2815 | 550 | 990 | 1540 |
| P36278 | NC_003896.1 | 2766 | Geminiviridae | ssDNA | 1 | 4 | 9 | 15 | 2766 | 440 | 990 | 1650 |
| P38608 | NC_003828.1 | 2773 | Geminiviridae | ssDNA | 1 | 5 | 9 | 14 | 2773 | 550 | 990 | 1540 |
| P27256 | NC_004005.1 | 2781 | Geminiviridae | ssDNA | 1 | 5 | 9 | 14 | 2781 | 550 | 990 | 1540 |
| P06946 | NC_003326.1 | 2750 | Geminiviridae | ssDNA | 1 | 5 | 9 | 11 | 2750 | 550 | 990 | 1210 |
| P0CK34 | NC_001439.1 (segment a) | 2647 | Geminiviridae | ssDNA | 1 | 5 | 8 | 12 | 2647 | 550 | 880 | 1320 |
| P27255 | NC_001934.1 (segment a) | 2593 | Geminiviridae | ssDNA | 1 | 5 | 8 | 12 | 2593 | 550 | 880 | 1320 |
| P27444 | NC_001936.1 (segment a) | 2633 | Geminiviridae | ssDNA | 1 | 5 | 8 | 12 | 2633 | 550 | 880 | 1320 |
| P03560 | NC_001507.1 (segment a) | 2588 | Geminiviridae | ssDNA | 1 | 5 | 8 | 11 | 2588 | 550 | 880 | 1210 |
| P36277 | NC_001938.1 (segment a) | 2601 | Geminiviridae | ssDNA | 1 | 5 | 8 | 12 | 2601 | 550 | 880 | 1320 |
| Q9DXE8 | NC_004044.1 | 2734 | Geminiviridae | ssDNA | 1 | 4 | 8 | 13 | 2734 | 440 | 880 | 1430 |
| P21942 | NC_001928.2 (segment a) | 2632 | Geminiviridae | ssDNA | 1 | 5 | 8 | 12 | 2632 | 550 | 880 | 1320 |
| P14966 | NC_001467.1 (segment a) | 2779 | Geminiviridae | ssDNA | 1 | 4 | 8 | 14 | 2779 | 440 | 880 | 1540 |
| Q96701 | NC_003866.1 (segment a) | 2583 | Geminiviridae | ssDNA | 1 | 5 | 8 | 12 | 2583 | 550 | 880 | 1320 |
| P29073 | NC_003379.1 | 2672 | Geminiviridae | ssDNA | 1 | 5 | 7 | 9 | 2672 | 550 | 770 | 990 |
| Q65386 | NC_003479.1 (segment 1) | 1043 | Nanoviridae | ssDNA | 1 | 7 | 11 | 12 | 1043 | 420 | 660 | 720 |
| Q9WIJ8 | NC_003559.1 (segment 10) | 999 | Nanoviridae | ssDNA | 1 | 5 | 9 | 10 | 999 | 300 | 540 | 600 |
| Q9Z035 | NC_003648.1 (segment 11) | 1001 | Nanoviridae | ssDNA | 1 | 6 | 9 | 10 | 1001 | 360 | 540 | 600 |
| P52501 | NC_001718.1 | 5075 | Parvoviridae | ssDNA | 1 | 4 | 11 | 13 | 5075 | 240 | 660 | 780 |
| P12930 | NC_001539.1 | 5323 | Parvoviridae | ssDNA | 1 | 5 | 10 | 13 | 5323 | 300 | 600 | 780 |
| P24840 | P24840 | 4938 | Parvoviridae | ssDNA | 1 | 5 | 10 | 13 | 4938 | 300 | 600 | 780 |
| P07297 | NC_001540.1 | 5517 | Parvoviridae | ssDNA | 1 | 4 | 9 | 12 | 5517 | 240 | 540 | 720 |
| P36310 | NC_001358.1 | 5176 | Parvoviridae | ssDNA | 1 | 5 | 9 | 12 | 5176 | 300 | 540 | 720 |
| P27404 | U13992.1 | 7677 | Caliciviridae | ssRNA | 3 | 3 | 5 | 6 | 7677 | 540 | 900 | 1080 |
| P27410 | NC_001543.1 | 7437 | Caliciviridae | ssRNA | 3 | 3 | 4 | 9 | 7437 | 540 | 720 | 1620 |
| Q04542 | L07418.1 | 7708 | Caliciviridae | ssRNA | 3 | 4 | 3 | 6 | 7708 | 720 | 540 | 1080 |
| P03608 | NC_004063.1 | 6318 | Tymoviridae | ssRNA | 3 | 1 | 2 | 2 | 6318 | 180 | 360 | 360 |
| P20123 | NC_001480.1 | 6331 | Tymoviridae | ssRNA | 3 | 3 | 4 | 4 | 6331 | 540 | 720 | 720 |
| P20124 | NC_001513.1 | 6211 | Tymoviridae | ssRNA | 3 | 3 | 3 | 2 | 6211 | 540 | 540 | 360 |
| Q82462 | NC_001981.1; NC_001982.1 | 7790 | Alphatetraviridae | ssRNA | 4 | 7 | 10 | 10 | 7790 | 1680 | 2400 | 2400 |
| Q9YRB2 | NC_001990.1 | 6625 | Alphatetraviridae | ssRNA | 4 | 8 | 11 | 14 | 6625 | 1920 | 2640 | 3360 |
| O12792 | PRJNA257543 | 6743 | Astroviridae | ssRNA | 3 | 5 | 11 | 19 | 6743 | 900 | 1980 | 3420 |
| Q3ZN05 | AY720891.1 | 6723 | Astroviridae | ssRNA | 3 | 5 | 11 | 20 | 6723 | 900 | 1980 | 3600 |
| Q4TWH7 | DQ028633.1 | 6762 | Astroviridae | ssRNA | 3 | 5 | 11 | 19 | 6762 | 900 | 1980 | 3420 |
| Q80KJ6 | NC_004579.1 | 6610 | Astroviridae | ssRNA | 3 | 5 | 10 | 15 | 6610 | 900 | 1800 | 2700 |
| P03591 | NC_001495.1 (segment 1) | 3644 | Bromoviridae | ssRNA | 3 | 4 | 9 | 11 | 3644 | 720 | 1620 | 1980 |
| P03601 | NC_003543.1 (segment 1) | 3171 | Bromoviridae | ssRNA | 3 | 6 | 9 | 11 | 3171 | 1080 | 1620 | 1980 |
| Q9DUT0 | NC_003649.1 (segment 1) | 3383 | Bromoviridae | ssRNA | 3 | 6 | 9 | 8 | 3383 | 1080 | 1620 | 1440 |
| P24402 | NC_004008.1 (segment 1) | 3158 | Bromoviridae | ssRNA | 3 | 5 | 8 | 10 | 3158 | 900 | 1440 | 1800 |
| P03602 | NC_002026.1 (segment 1) | 3234 | Bromoviridae | ssRNA | 3 | 5 | 8 | 9 | 3234 | 900 | 1440 | 1620 |
| P69466 | NC_002034.1 (segment 1) | 3357 | Bromoviridae | ssRNA | 3 | 7 | 8 | 12 | 3357 | 1260 | 1440 | 2160 |

| | | | | | | | | | | | | |
|---|---|---|---|---|---|---|---|---|---|---|---|---|
| Q9YNE2 | NC_003673.1 (RNA1) | 3128 | Bromoviridae | ssRNA | 3 | 7 | 7 | 9 | 3128 | 1260 | 1260 | 1620 |
| P03598 | NC_003844.1 (segment 1) | 3491 | Bromoviridae | ssRNA | 3 | 5 | 7 | 12 | 3491 | 900 | 1260 | 2160 |
| P23627 | NC_003837.1 (segment 1) | 3410 | Bromoviridae | ssRNA | 3 | 6 | 7 | 11 | 3410 | 1080 | 1260 | 1980 |
| Q03500 | NC_001434.1 | 7176 | Hepeviridae | ssRNA | 3 | 5 | 7 | 7 | 7176 | 900 | 1260 | 1260 |
| P29326 | M73218.1 | 7207 | Hepeviridae | ssRNA | 3 | 5 | 7 | 7 | 7207 | 900 | 1260 | 1260 |
| P33426 | AF444002.1 | 7204 | Hepeviridae | ssRNA | 3 | 5 | 7 | 7 | 7204 | 900 | 1260 | 1260 |
| Q04611 | D10330.1 | 7194 | Hepeviridae | ssRNA | 3 | 5 | 7 | 7 | 7194 | 900 | 1260 | 1260 |
| P03612 | NC_001417.2 | 3569 | Leviviridae | ssRNA | 3 | 3 | 6 | 5 | 3569 | 534 | 1068 | 890 |
| P03614 | X15031.1 | 3575 | Leviviridae | ssRNA | 3 | 3 | 6 | 5 | 3575 | 534 | 1068 | 890 |
| P07234 | X03869.1 | 3466 | Leviviridae | ssRNA | 3 | 3 | 6 | 5 | 3466 | 534 | 1068 | 890 |
| P09510 | NC_004750.1 | 5677 | Luteoviridae | ssRNA | 3 | 7 | 14 | 20 | 5677 | 1260 | 2520 | 3600 |
| Q84711 | NC_003629.1 | 5705 | Luteoviridae | ssRNA | 3 | 7 | 13 | 16 | 5705 | 1260 | 2340 | 2880 |
| P17522 | NC_001747.1 | 5987 | Luteoviridae | ssRNA | 3 | 7 | 13 | 22 | 5987 | 1260 | 2340 | 3960 |
| P09514 | NC_003743.1 | 5641 | Luteoviridae | ssRNA | 3 | 7 | 12 | 21 | 5641 | 1260 | 2160 | 3780 |
| P12871 | NC_002690.1; NC_002691 | 4540 | Nodaviridae | ssRNA | 3 | 8 | 13 | 18 | 4540 | 1440 | 2340 | 3240 |
| Q91D83 | NC_003448.1; NC_003449.1 | 4528 | Nodaviridae | ssRNA | 3 | 6 | 11 | 13 | 4528 | 1080 | 1980 | 2340 |
| P36290 (VP1, VP2, VP3, VP4) | D90457 | 7461 | Picornaviridae | ssRNA | 3 | 4, 3, 2, 1 | 9, 4, 1, 2 | 7, 5, 3, 0 | 7461 | 600 | 960 | 900 |
| P12915 (VP1, VP2, VP3, VP4) | NC_001859.1 | 7414 | Picornaviridae | ssRNA | 3 | 4, 2, 2, 1 | 9, 2, 2, 1 | 9, 4, 1, 1 | 7414 | 540 | 840 | 900 |
| P32537 (VP1, VP2, VP3, VP4) | NC_030454.1 | 7313 | Picornaviridae | ssRNA | 3 | 4, 4, 3, 2 | 6, 3, 2, 1 | 9, 5,3, 0 | 7313 | 780 | 720 | 1020 |
| Q82122 (VP1, VP2, VP3, VP4) | L24917 | 7124 | Picornaviridae | ssRNA | 3 | 3, 2, 4, 1 | 8, 3, 5, 2 | 9, 3, 4, 0 | 7124 | 600 | 1080 | 960 |
| O91734 (VP1, VP2, VP3, VP4) | AF029859 | 7397 | Picornaviridae | ssRNA | 3 | 5, 2, 3, 1 | 7, 2, 4,1 | 8, 1, 4, 0 | 7397 | 660 | 780 | 780 |
| P03304 (VP1, VP2, VP3, VP4) | NC_001479.1 | 7835 | Picornaviridae | ssRNA | 3 | 4, 4, 2, 1 | 6, 6, 4, 0 | 6, 11, 6, 0 | 7835 | 660 | 1020 | 1380 |
| O73556 (VP1, VP2, VP3, VP4) | NC_038319.1 | 7339 | Picornaviridae | ssRNA | 3 | 3,4,4,1 | 4, 5, 7, 0 | 4, 9, 6, 0 | 7339 | 660 | 960 | 1140 |
| Q05057 (CP1,CP2,CP3) | NC_003628.1 | 9871 | Secoviridae | ssRNA | 3 | 2, 4, 4 | 3, 5, 8 | 3, 3, 9 | 9871 | 600 | 960 | 900 |
| Q83034 (CP1,CP2,CP3) | NC_001632.1 | 12226 | Secoviridae | ssRNA | 3 | 3, 2, 5 | 3, 4, 8 | 2, 6, 9 | 12226 | 600 | 900 | 1020 |
| Q9DSN8 (VP1, VP2, VP3, VP4) | NC_002548.1 | 9491 | Dcistroviridae | ssRNA | 3 | 3, 2, 1, 4 | 3, 2, 3, 7 | 2, 3, 5, 7 | 9491 | 600 | 900 | 1100 |
| P13418 (VP1, VP2, VP3, VP4) | NC_003924.1 | 9185 | Dicistroviridae | ssRNA | 3 | 4, 1, 3, 3 | 0, 1, 2, 7 | 6, 0, 8, 6 | 9185 | 660 | 600 | 1200 |
| P13897 | NC_003908.1 | 11484 | Togaviridae | ssRNA | 4 | 6 | 16 | 23 | 11484 | 1440 | 3840 | 5520 |
| P89946 | NC_001786.1 | 11486 | Togaviridae | ssRNA | 4 | 6 | 15 | 22 | 11486 | 1440 | 3600 | 5280 |
| Q306W5 | NC_003899.1 | 11675 | Togaviridae | ssRNA | 4 | 6 | 15 | 23 | 11675 | 1440 | 3600 | 5520 |
| P03315 | NC_003215.1 | 11442 | Togaviridae | ssRNA | 4 | 6 | 14 | 23 | 11442 | 1440 | 3360 | 5520 |
| Q5XXP3 | NC_004162.2 | 11826 | Togaviridae | ssRNA | 4 | 7 | 13 | 23 | 11826 | 1680 | 3120 | 5520 |
| Q8JJX0 | NC_003930.1 | 11918 | Togaviridae | ssRNA | 4 | 6 | 13 | 21 | 11918 | 1440 | 3120 | 5040 |
| Q8QL52 | NC_003433.1 | 11900 | Togaviridae | ssRNA | 4 | 6 | 13 | 22 | 11900 | 1440 | 3120 | 5280 |
| P36329 | NC_001449.1 | 11444 | Togaviridae | ssRNA | 4 | 7 | 13 | 23 | 11444 | 1680 | 3120 | 5520 |
| Q5Y388 | NC_006558.1 | 11597 | Togaviridae | ssRNA | 4 | 6 | 12 | 22 | 11597 | 1440 | 2880 | 5280 |
| Q9JGK8 | AB032553.1 | 11698 | Togaviridae | ssRNA | 4 | 6 | 12 | 22 | 11698 | 1440 | 2880 | 5280 |
| P22056 | NC_001512.1 | 11835 | Togaviridae | ssRNA | 4 | 7 | 12 | 22 | 11835 | 1680 | 2880 | 5280 |
| Q6X2U3 | NC_001545.2 | 9762 | Togaviridae | ssRNA | 4 | 6 | 8 | 12 | 9762 | 1440 | 1920 | 2880 |
| P89036 | NC_002598.1 | 4326 | Tombusviridae | ssRNA | 3 | 5 | 10 | 13 | 4326 | 900 | 1800 | 2340 |
| P22955 | NC_003756.1 (segment 1) | 3890 | Tombusviridae | ssRNA | 3 | 5 | 9 | 12 | 3890 | 900 | 1620 | 2160 |
| P11689 | NC_001554.1 | 4775 | Tombusviridae | ssRNA | 3 | 6 | 8 | 13 | 4775 | 1080 | 1440 | 2340 |
| Q66098 | NC_003530.1 (segment 1) | 3840 | Tombusviridae | ssRNA | 3 | 4 | 9 | 13 | 3840 | 720 | 1620 | 2340 |
| P22959 | NC_001777.1 | 3680 | Tombusviridae | ssRNA | 3 | 5 | 9 | 13 | 3680 | 900 | 1620 | 2340 |
| P11642 | NC_003627.1 | 4437 | Tombusviridae | ssRNA | 3 | 5 | 9 | 10 | 4437 | 900 | 1620 | 1800 |
| P15183 | NC_001469.1 | 4699 | Tombusviridae | ssRNA | 3 | 5 | 7 | 13 | 4699 | 900 | 1260 | 2340 |
| P17456 | NC_003532.1 | 4732 | Tombusviridae | ssRNA | 3 | 5 | 7 | 13 | 4732 | 900 | 1260 | 2340 |
| Q9QBU3 | NC_000939.2 | 4415 | Tombusviridae | ssRNA | 3 | 6 | 7 | 12 | 4415 | 1080 | 1260 | 2160 |

Geminiviruses usually formed by two fusioned T=1 capsid, totalizing 110 subunits.
Pseudo T=3, with 60 subunits of each protein derived from polyprotein.
Pseudo T=7, 72 pentamers of L1/VP1 + 72 copies of L2/VP2
Pseudo T=3, 178 VP1 subunits

### Supplemental information S2

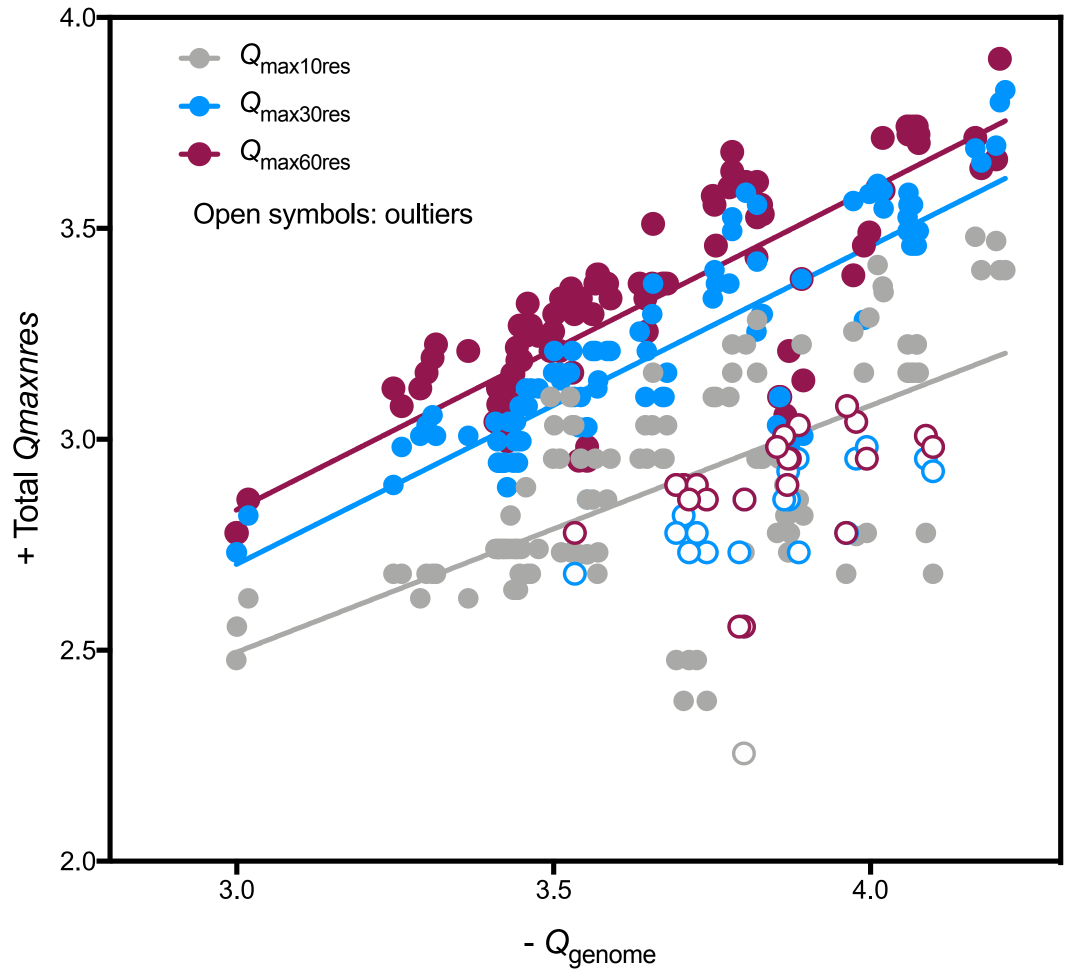

**Legend S2:** The maximum net-charge value found in 10, 30 or 60 amino acid residues stretch was multiplied by the number of subunits forming the capsid (Total $Q_{max}$) of 129 viruses from 25 different families. The total nucleic-acid net charge was calculated from the number of nucleotides residues in the genome ($Q_{genome}$). For multipartide viruses, the longest genome segment was considered for the plot. A straight line fit for the entire dataset was calculated and a shaded area indicates the outliers. Complete statistics for the linear fit are shown below:

| | 10 | 30 | 60 |
|---|---|---|---|
| Straight line | | | |
| Best-fit values | | | |
| YIntercept | 0.7416 | 0.4417 | 0.5524 |
| Slope | 0.5846 | 0.7541 | 0.7602 |
| Std. Error | | | |
| YIntercept | 0.2256 | 0.1586 | 0.1862 |
| Slope | 0.06095 | 0.0432 | 0.05074 |
| 95% CI (profile likelihood) | | | |
| YIntercept | 0.2952 to 1.188 | 0.1273 to 0.7562 | 0.1833 to 0.9216 |
| Slope | 0.464 to 0.7053 | 0.6684 to 0.8397 | 0.6596 to 0.8608 |
| Goodness of Fit | | | |
| Degrees of Freedom | 126 | 106 | 105 |
| R square | 0.422 | 0.7419 | 0.6813 |
| Absolute Sum of Squares | 4.408 | 1.665 | 2.265 |
| Sy.x | 0.187 | 0.1253 | 0.1469 |
| Number of points | | | |
| # of X values | 129 | 129 | 129 |
| # Y values analyzed | 129 | 129 | 129 |
| Outliers (excluded, Q=2%) | 1 | 21 | 22 |

#### Supplemental information S3

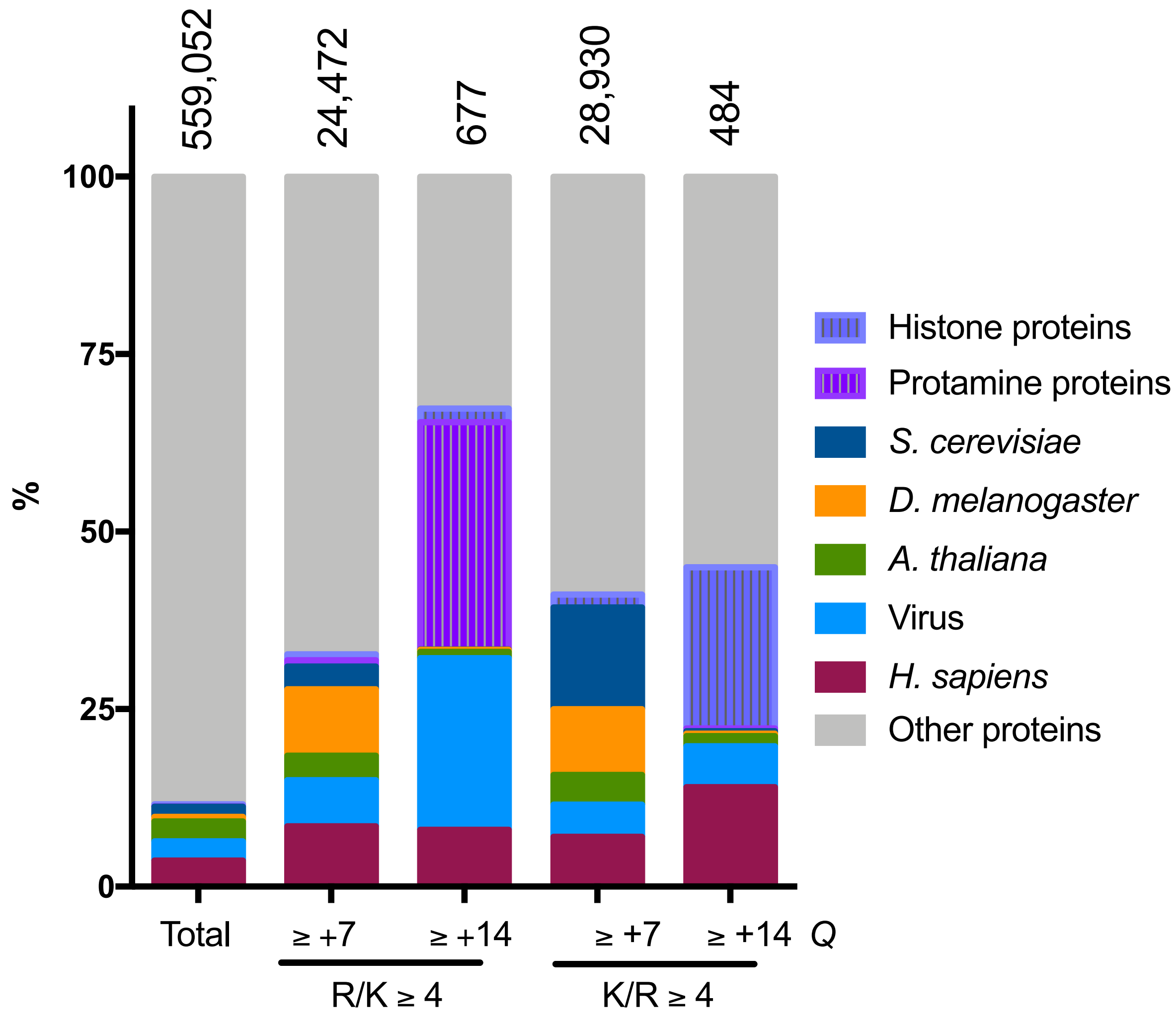

Figure S3: Proportion of proteins containing positively charged segments in the Swiss-Prot data base. Protein sequences derived from the reviewed Swiss-Prot data- bank were used as input for a program that calculates the net charge of every consecutive 30 residue amino-acid segments ($Q_{30res}$). The arginine and lysine residues of fragments with $Q_{30res} \geq 7$ or + ≥ 14 were determined. Proteins containing at least one segment with $Q_{30res}$ ≥ +7 or ≥ +14 with R/K ≥ 4 were listed according to the organism or function.
